# Redox Regulation in O_2_-Tolerant [FeFe] Hydrogenases: Insights from two homologues

**DOI:** 10.64898/2026.05.07.723305

**Authors:** Rinat Khundoker, Sean H. Majer, Alexey Silakov

**Author notes:** Corresponding author: Alexey Silakov,.

## Abstract

O_2_-tolerance is a desirable property for [FeFe] hydrogenases, which are highly efficient H_2_-producing catalysts. While most such enzymes are highly sensitive to aerobic environments, a small number of explored representatives exhibit exceptional stability and even H_2_-producing activity under oxygenic conditions. However, the genetic signatures of the O_2_-tolerance in this class of enzymes remain largely unknown. To address this knowledge gap, we explored a close homologue of a well-characterized O_2_-tolerant [FeFe] hydrogenase from *Clostridium beijerinckii* (*Cb*HydA1) - a hydrogenase from *Terrisporobacter glycolicus* (*Tg*HydA1). Our investigation indeed confirms that *Tg*HydA1 can transition to the O_2_-stable H_inact_ state, a hallmark of O_2_ tolerance. The surprising outcome is that despite the high amino acid similarity, *Tg*HydA1 shows a substantially higher propensity to remain in the H_inact_ state than *Cb*HydA1. Using protein film electrochemical experiments, we demonstrate that the root of this behavior lies in roughly tenfold slower reactivation rates than those of *Cb*HydA1 at any applied potential. This degree and direction of variation in reactivation kinetics have not been observed before for any other O_2_-tolerant [FeFe] hydrogenases or their variants to date, uncovering a yet-to-be-explored facet of reactivity alteration available to these enzymes. Overall, the results presented here highlight the importance of a holistic analysis of [FeFe] hydrogenase sequences in the context of their interaction with O_2_ that encompasses the protein environment and properties of the auxiliary metallocofactors.

## INTRODUCTION

Hydrogenases represent a structurally diverse class of metalloenzymes that can catalyze the reversible oxidation of dihydrogen.[1–4] [FeFe] hydrogenases comprise a family of such enzymes that contain a complex six-Fe active cofactor called the H-cluster (see **Fig. 1**).[1, 5–7] These enzymes are of particular interest to industrial applications in H_2_-producing biological and bio-hybrid systems.[8–11]

**Figure 1.**
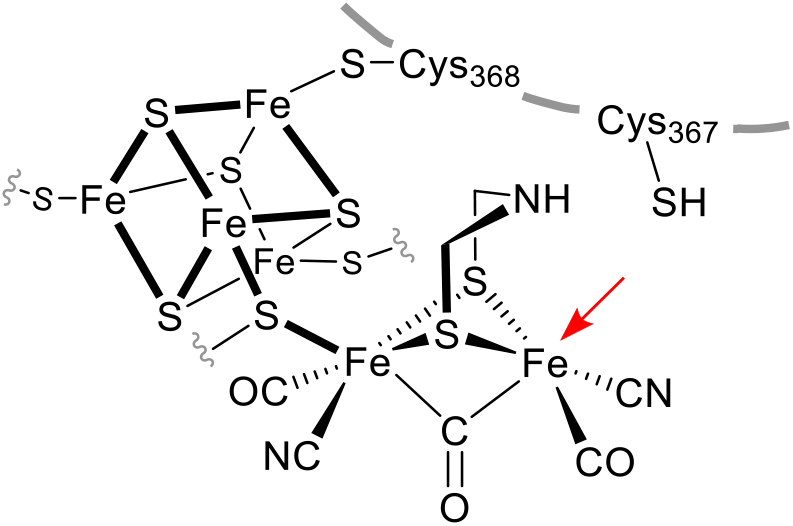
Schematic representation of the active site of [FeFe] hydrogenases, the H-cluster. The red arrow indicates the open coordination site. The Cys residue numbering is according to the *Cb*HydA1 amino acid sequence.

Unfortunately, the vast majority of known [FeFe] hydrogenases can only operate under strictly anaerobic conditions, significantly limiting their industrial use.[1, 2, 12] However, recent discoveries of a group A [FeFe] hydrogenase from *Clostridium beijerinckii* (*Cb*HydA1) that shows enhanced stability in the presence of O_2_ renew interest in this family of enzymes as potential H_2_ catalysts in renewable energy technology for H_2_ production.[13–16] The key feature of *Cb*HydA1 is its ability to rapidly and reversibly access an O_2_-stable, inactive state, called H_inact_, under oxidizing conditions. Our research demonstrated that this state is one electron more oxidized than the active resting state H_ox_ (see Scheme S1 in Supporting information).[14] Past research by us [14, 17], and others [7, 18–20] concluded that a highly conserved proton channeling Cys residue (Cys367 in *Cb*HydA1) near the H-cluster plays a key role in the stabilization of H_inact_ (see **Fig. 1**). Our recent coupling of *Cb*HydA1 to a PsaE-deleted cyanobacterial photosystem I (PSI_ΔPsaE_) by engineering a *Cb*HydA1-PsaE fusion showed the ability of the PSI_ΔPsaE_:*Cb*HydA1-PsaE nanoconstruct to produce H_2_ under fully aerobic conditions.[21] This result is of particular importance as it demonstrates the ability of *Cb*HydA1 to operate on air, i.e., it exhibits some degree of O_2_-tolerance. Thus, we can apply this term the same way it has been used to designate a class of O_2_-tolerant [NiFe] hydrogenases, which rapidly interconvert between the active state and the “ready” inactive Ni(III)-Fe(II) state Ni-B without fully inactivating into a Ni-A state. [22–24] For completeness, we note that a subclass of [NiFeSe] hydrogenases can maintain O_2_-tolerance without interconverting to the “overoxidized” Ni(III) species (Ni-B or Ni-A).[25–28]

A more recent discovery of an O_2_-tolerant group B [FeFe] hydrogenase demonstrates that this property is not restricted to group A *Cb*HydA1-homologues.[29] We also demonstrated further expansion of [FeFe] hydrogenase repertoire by demonstrating the existence of an H_2_-dependent H_2_O_2_ reductase in *Clostridium perfringens* (*Cper*HydR) that utilizes an H-cluster to generate reducing equivalents from H_2_ for the reduction of H_2_O_2_ by a coexisting rubrerythrin subdomain of the same protein. In our in vitro experiments, *Cper*HydR was able to withstand the presence of high concentrations of H_2_O_2_ without losing activity. Overall, these studies indicate that tolerance to strong oxidizing agents by [FeFe] hydrogenases is far more widespread in nature than previously thought.[1, 30, 31] Then the question is, how can we recognize such a desirable trait?

Unfortunately, the genetic signatures that hallmark the O_2_ tolerance of [FeFe] hydrogenases have yet to be resolved, presenting a barrier to identifying similar, better-performing enzymes for further optimization of bioengineered H_2_-producing systems. We proposed that increased protein flexibility allows Cys367 of *Cb*HydA1 to adopt rotamers inaccessible to the O_2_-sensitive enzymes.[14, 17] Therefore, the relationship between the ability of these [FeFe] hydrogenases to safely inactivate and the amino acid sequence is expected to be complex. Indeed, site-specific amino acid substitutions on the protein surface affected O_2_-tolerance in *Cb*A5H, a near-identical homologue of *Cb*HydA1.[20] *Cb*HydA1-PsaE fusion showed altered oxidative inactivation behavior in protein film voltammetry (PFV) experiments, emphasizing the importance of the overall protein dynamic.[21] Molecular dynamics simulations also suggested the relevance of another set of hydrophobic residues near Cys367 to the O_2_-tolerance of [FeFe] hydrogenases.[29] Overall, these past studies bring to light the challenge in deciphering the genetic signatures of O_2_-tolerance, as it cannot be attributed to a small subset of amino acid alterations.

To understand the degree of sequence similarity required to retain O_2_-tolerance, our laboratory is interested in investigating close homologues of *Cb*HydA1. In this manuscript, we present a case of a group A [FeFe] hydrogenase from *Terrisporobacter (T*.*) glycolicus* strain KPPR-9 (*Tg*HydA1). *T. glycolicus* (previously known as *Clostridium glycolicum*) is an obligatory anaerobic, spore-forming, Gram-positive, rod-shaped bacterium that was first isolated from mud as an anaerobe utilizing ethylene glycol as a carbon source.[32, 33] Kusel et al. showed that RD-1, a strain of *T. glycolicus* extracted from the root of sea grass *Halodule wrightii*, is an acetogen and can ferment fructose, glucose, maltose, and xylose into acetate and ethanol.[34] This organism can produce H_2_ during fermentation. *T. glycolicus* genome contains two different enzymes annotated as [FeFe]-hydrogenases, one of which is highly homologous to *Cb*HydA1 (Locus tag: SAMN02910355_3663, NCBI Code: SFJ67572.1, we term *Tg*HydA1) and is the focus of this study. The second putative [FeFe] hydrogenase (Locus tag: SAMN02910355_3560, NCBI Code: SFJ66003.1, we term *Tg*HydA2) is likely of M2 type, i.e., containing two accessory metallocofactors. It harbors “TTLCP” variation of the L2 conserved sequence instead of the typical group A “TSCCP” code, i.e., lacking the conserved proton-channeling Cys required for bidirectional hydrogenase activity. We show that *Tg*HydA1 exhibits a surprising divergence in the inactivation process despite high sequence similarity with *Cb*HydA1 (84.21%, **Fig. S1**). Using electrochemical assays, we show that the phenomenological origin of this discrepancy lies in a substantially altered energetics of the reactivation process, poising this enzyme towards H_inact_ state under typical laboratory *in vitro* conditions.

## RESULTS AND DISCUSSIONS

We cloned *Tg*HydA1 into the pET28a vector and transformed it into *E. coli* BL21(DE3) ΔiscR cells for heterologous protein production. We augmented the gene with appropriate affinity tags to facilitate purification, a method we successfully used previously to produce *Cb*HydA1. After isolation, we reconstituted the FeS clusters and incorporated the H-cluster using a synthetic [2Fe] precursor using well established procedures.[14, 21, 35]

We noticed that after a typical sample preparation (see supporting information), *Tg*HydA1 is consistently found in the inactive state (H_inact_), despite employing strict anaerobic conditions and requiring an excess of strong reductants during metallocofactor incorporation steps (see **Fig. 2A**). In our previous work, similar sample preparation procedures yielded fully active *Cb*HydA1.[14, 17] We note that while the fourier-transform infrared (FTIR) spectra of the H_inact_ state are similar between *Cb*HydA1 and *Tg*HydA1, there are noticeable differences in the exact positions of the CO bands. These discrepancies indicate small but detectable variation in the electronic structure of the H-cluster (see **Table S1**). Surprisingly, we were unable to obtain IR spectra of the pure active states at room temperature by incubating *Tg*HydA1 with a strong reductant, such as sodium dithionite (DT; **Fig. 2B**). To test whether the protein requires a stronger reductant for reactivation, we used Ti(III)-citrate. The results were very similar (**Fig. 2C**), i.e., the sample largely remained in the H_inact_ state. We noted, however, the appearance of additional unidentified bands that somewhat increased in intensity between DT- and Ti(III)-citrate-treated samples. Based on the IR bands’ positions, we preemptively assign them to the H_trans_ state as previously observed in *Dd*HydAB (*Dd*=*Desulfovibrio desulfuricans*), [36] although we cannot fully exclude the H_hyd_(H^+^) assignment either. A further investigation is needed to confirm this hypothesis, which is beyond the scope of this work.

**Figure 2.**
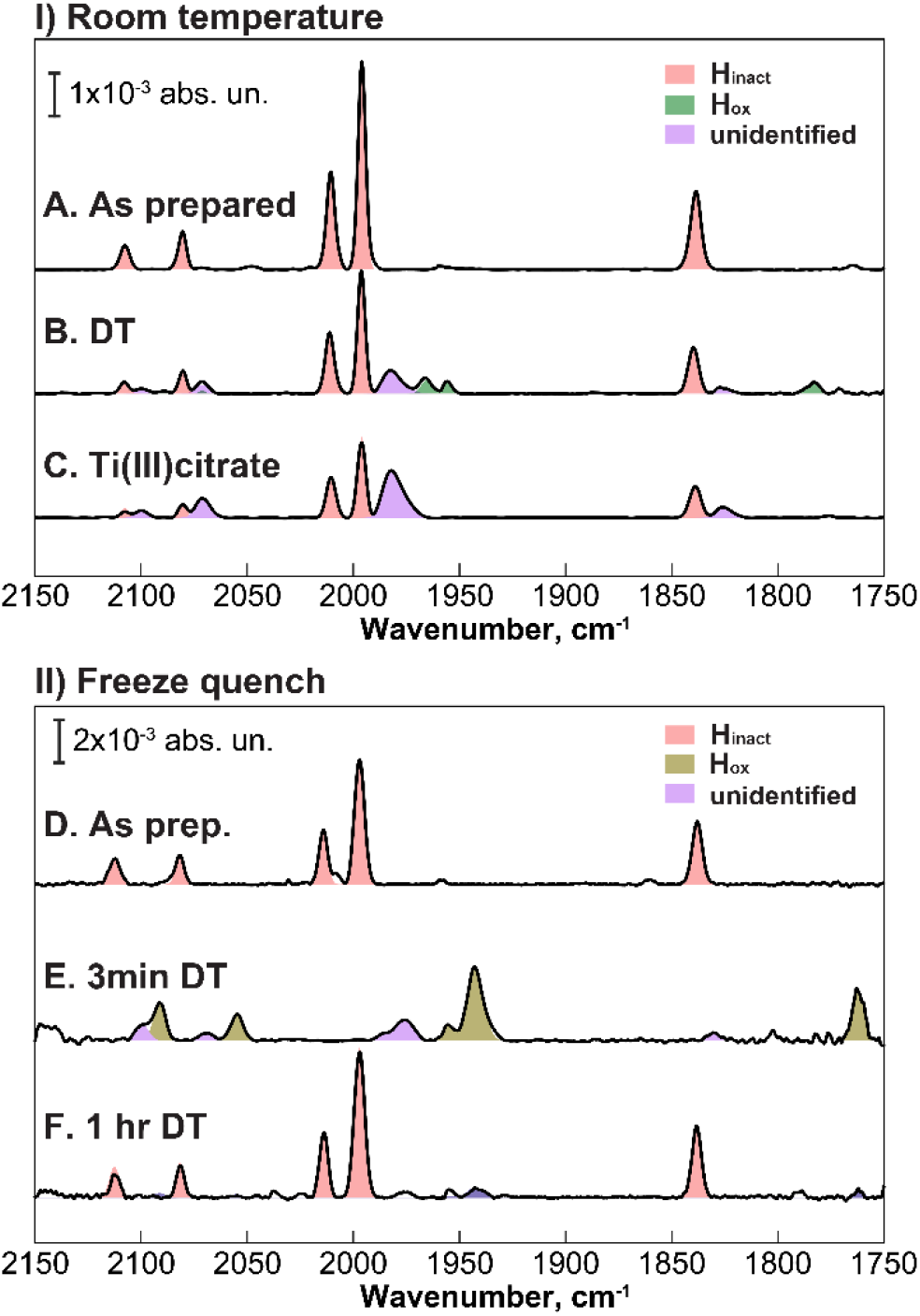
FTIR spectra of *Tg*HydA1 obtained at room temperature (panel I) and at 15 K (panel II). A,D) as-prepared samples. B,C) As-prepared *Tg*HydA1 exposed to 5 mM sodium dithionite (DT) and 5 mM Ti(III) citrate, respectively. E,F) freeze-quenching reduction with 5 mM DT after 3 min and 1 hr, respectively.

Since *Tg*HydA1 shows a reasonable H_2_ evolution rate of 136±24 µmol(H_2_) mg(prot)^-1^ min^-1^ (see below), we are inclined to think that the enzyme self-inactivates within the timeframe required to load the sample into the IR cell and record FTIR spectra of sufficient quality (ca. 30 min). Given the high enzyme concentrations required for FTIR, it is possible that the reductant may couple effectively with H_2_ evolution, leading to auto-oxidation of *Tg*HydA1 during a room-temperature measurement. Unfortunately, measuring a low-concentration protein sample in our FTIR spectrometer is impractical. Therefore, to test this hypothesis, we performed a freeze-quench experiment in which we froze the *Tg*HydA1 sample shortly after adding sodium dithionite. Indeed, freezing the DT-treated sample immediately after preparation (within 3 min) afforded detection of the active H_ox_ state by a cryogenic FTIR at 15 K (**Fig. 2E**). A similar experiment in which we waited 1 hour before the freeze-quench showed a complete conversion of the enzyme back to the H_inact_ state despite maintaining a strictly anaerobic (3% H_2_, 97%N) atmosphere (see **Fig. 2F**).

These experiments reassured us that *Tg*HydA1 is enzymatically active but tends to remain in the H_inact_ state in the absence of a strong reducing environment. This is a substantial departure from the typical behavior of *Cb*HydA1, which remains active unless exposed to an oxidant or placed in a low-pH environment (below pH 6.5).[14, 17] Hence, we attribute this effect to differences in the inactivation and reactivation kinetics.

To further investigate this behavior, we performed a series of protein film voltammetry experiments at various pH. The H^+^ reduction (negative) current dominates the acquired cyclic voltammetry (CV) traces even at high pH, where [FeFe] hydrogenases should be thermodynamically biased towards H_2_ oxidation (positive current),[37–40] see **Fig. S2**. Another unusual feature of the traces observed was the presence of substantial hysteresis around the H^+^/H_2_ thermodynamic equilibrium potential between forward and reverse potential scans, most prominent at higher scan rates (100 mV/s, see **Fig. 3**). When compared to CV traces of *Cb*HydA1 measured at identical conditions, the inhibition of the H_2_-oxidation current due to inactivation occurs at less oxidizing potential than *Cb*HydA1, concurring with the observed higher tendency of *Tg*HydA1 to inactivate when presented with oxidizing environment (above E_0_(H^+^/H_2_)).

**Figure 3.**
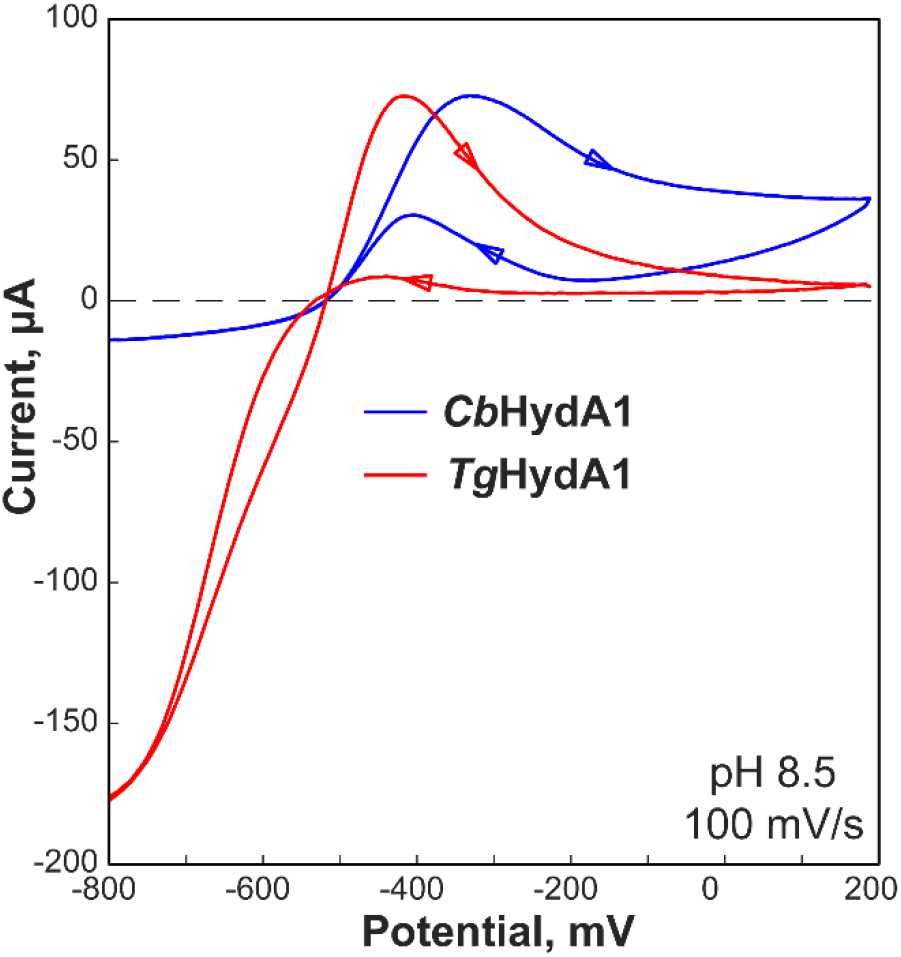
Comparison of protein film cyclic voltammetry traces of *Cb*HydA1 and *Tg*HydA1 performed at pH 8.5 and with 100 mV/s scan rate. The trace of *Cb*HydA1 was scaled up by a factor of 2 to match the maximum of the oxidizing current of *Tg*HydA1. Experiments were performed at room temperature under 1 atm H_2_.

To rationalize the kinetics of inactivation, we conducted a series of chronoamperometry experiments with *Cb*HydA1 and *Tg*HydA1 involving repeated steps between more oxidizing (inactivating) and more reducing (activating) potentials starting with a fully activating potential (-800 mV vs NHE, see **Fig. S3** and **Fig. S4**).[40, 41] The analysis of the catalytic current progression after a potential step allows us to extract kinetic rates related to interconversion between the active and inactive states. Following the work of Winkler et al.,[20] we adopted a model assuming the presence of two active states in a dynamic equilibrium:

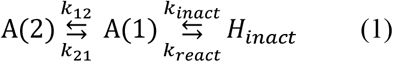

Our analysis of the heterogeneous EPR spectra of the H_ox_ and H_ox_-CO states of *Cb*HydA1 concluded that the H-cluster exhibits two spectroscopically distinct structural isoforms in the active state, fully consistent with the aforementioned kinetic model.[17] While analytical equations for such a process are trivial to obtain, the dependence of the exponential and pre-exponential factors on the kinetic constants is highly complex (see Supporting Information). Thus, fitting the chronoamperometry data to kinetic constants is an ill-posed problem that requires regularization assumptions to yield meaningful results. Winkler et al.[20] postulated that the rates k_21_ and k_12_ are independent of potential, consistent with our spectroscopic findings of two structural isoforms in dynamic equilibrium.[17]

Considering that the Cys367 rearrangement is likely the rate-limiting step in the oxidative inactivation, the transition from the H_ox_ state to the H_inact_ state should have little-to-no potential dependence, leading to apparent potential independence of k_inact_. Then the potential dependence of the kinetic processes derives solely from the rate of reduction of the H_inact_ state (k_react_), i.e. it depends on the potential-dependent electron-transfer rates from the H-cluster as governed by the redox properties of the H-cluster in the H_inact_ state and the accessory metallocofactors. **Fig. 4** displays the results of potential-step chronoamperometry fits performed at pH 8.5 with the model described above. We have observed very similar kinetic rates for k_12_ and k_21_ between *Cb*HydA1 and *Tg*HydA1. Since these rates are associated with stochastic conversion between two active isoforms, the similarity between two enzymes indicates a similarity of their structural elasticity. The similarity in the k_inact_ rates between the two species further supports this conclusion.

**Figure 4.**
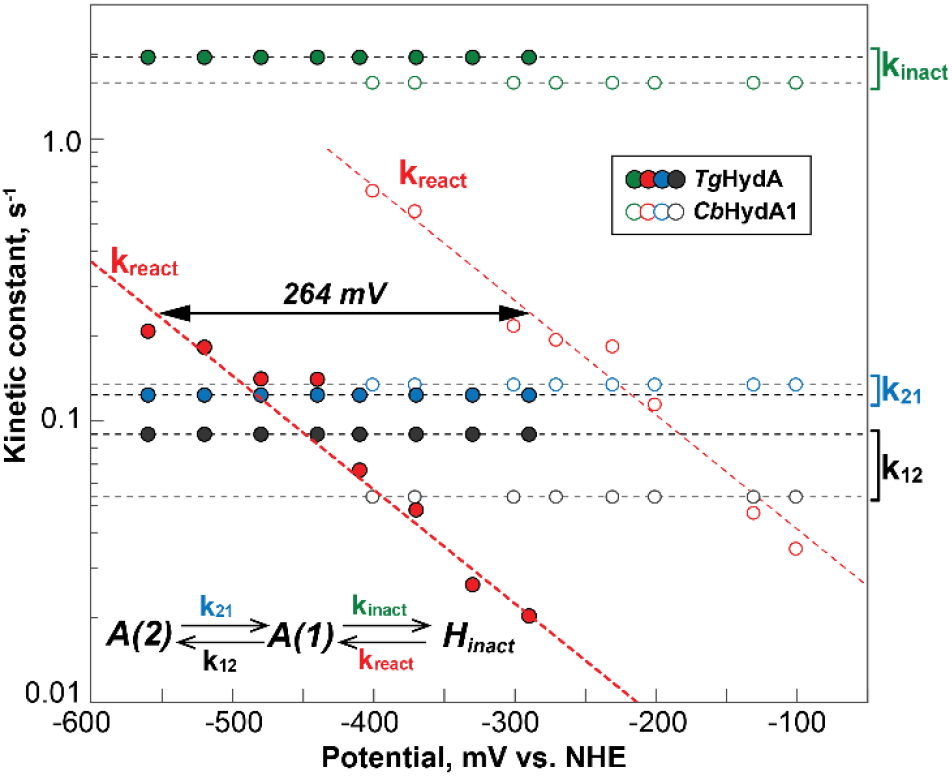
Potential dependence of kinetic constants extracted from chronoamperometry experiments of *Tg*HydA1(filled circles) and *Cb*HydA1 (hollow circles) using an AAI model described in the main text and schematically depicted in the bottom left corner of the plot.

The most notable result of this analysis is that the k_react_ values are universally about an order of magnitude *lower* for *Tg*HydA1 than those of *Cb*HydA1 at any given potential. This observation phenomenologically explains the propensity of *Tg*HydA1 to remain in the H_inact_ state under conditions that normally afford us accumulation of active states in other [FeFe] hydrogenases.[14, 17, 37]

As to the underlying cause, past precedents have demonstrated that even amino acid modifications away from the active site can influence the O_2_-tolerance.[18, 21] However, no reported amino acid substitutions have resulted in a *decrease* in the reactivation rate, as we have observed here. Furthermore, all previously explored amino acid motifs in *Cb*HydA1 are completely conserved in *Tg*HydA1 (see **Fig. S1**). It is thus likely that there is no significant alteration to the structural mobility between two proteins around the active site, as indeed suggested by the similarity of k_12_, k_21_, and k_inact_ rates. We thus propose that the effect originates largely from the differences in the redox potentials of the metallocofactors, H-cluster, and the accessory [4Fe-4S] clusters. Given the dramatic change in the k_react_(E) dependence and the observation of an H_trans_-like state in IR, we incline to suggest that the modification of the H-cluster’s redox potential of inactivation and reactivation roots in an alteration away from the motifs explored so far. However, further investigation into the electronic properties of the metallocofactors of *Cb*HydA1 and *Tg*HydA1 is needed to clarify this point.

Our earlier work demonstrated that once in the H_inact_ state, the *Cb*HydA1 remained fairly stable for a long time under fully aerobic conditions (<20% degradation after 44 hours at room temperature).[14] However, we and others have noted a gradual decrease in *Cb*HydA1 activity upon repeated cycles of exposure to O_2_ and reactivation.[14, 18] It is thus likely that the protein’s transition into the inactive state is not fast enough to avoid harmful ROS-generating O_2_ reduction. Given the increased propensity to remain in the H_inact_ state, we hypothesized that *Tg*HydA1 may exhibit less O_2_-induced degradation when reacted with oxygen. To test this idea, we repeatedly exposed low-concentration protein samples (1 μM) to air at 4°C for 70 min in the absence of any oxygen-scavenging agents such as sodium dithionite. Prior to each exposure, the enzyme was extensively activated under 1 atm H_2_ for 70 min. Subsequently, we analyzed the H_2_-evolving activity of diluted sample aliquots (at 20 nM concentration) by gas chromatography (see Materials and Methods). To separate general protein degradation during prolonged sample treatments from the effect of exposure to O_2_, we have performed parallel experiments in which aliquots of the same sample remained inside the anaerobic chamber (3%H_2_, 97%N_2_) but were otherwise treated identically. Notably, we observed a substantial protein degradation at pH below 7.0, even in the absence of any sample treatment (see **Fig. S6**). Therefore, we performed the experiments shown here at somewhat higher pH of 7.5 that allowed to considerably stabilize *Tg*HydA1. **Fig. 5** shows the comparison of the H_2_ evolution activity of aerobically and anaerobically treated (control) samples.

**Figure 5.**
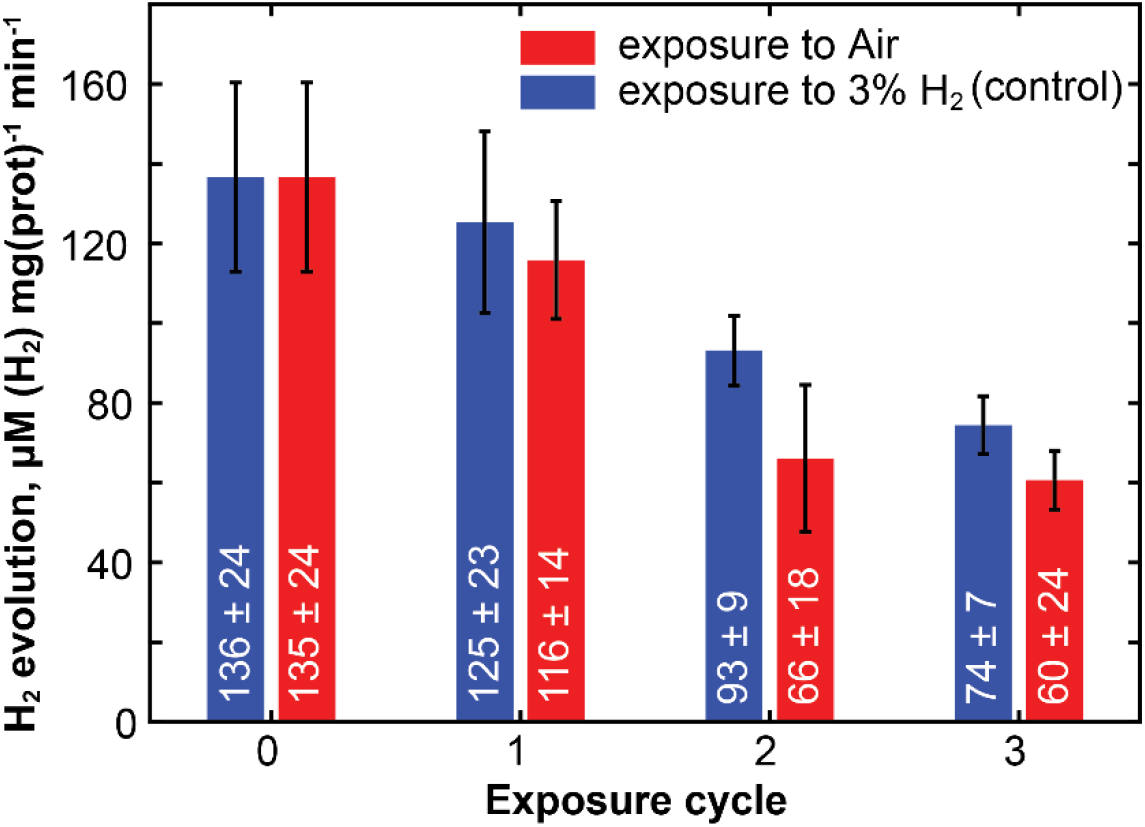
H_2_ evolution activity of *Tg*HydA1 after repeated expose to air at pH 7.5. Prior to each exposure, a 1 μM protein solution was activated by incubating under 1 atm H_2_ for 70 min. Subsequently, samples were either taken out the anaerobic chamber for 70 min on ice (red bars) or left in the chamber for the same period (blue bars). Activity assays were performed by gas chromatography. Error bars are standard deviations calculated from measurements in triplicate.

The key result of this experiment is that, even in the absence of sodium dithionite, exposure of TgHydA1 to air results in only a minor loss of activity (<20% after three cycles). Therefore, it appears that an increased propensity for inactivation in *Tg*HydA1 indeed coincides with protein stability, at least with respect to the interaction with O_2_. This trend is encouraging and warrants further investigation into why this protein demonstrated greater stability upon aerobic inactivation and whether this translates into enhanced O_2_ tolerance. It would be interesting to see how this protein performs under aerobic conditions when coupled to a photosystem I complex; a study planned for our laboratory in the future.

Lastly, we would like to point out that acetogens, such as *T. glycolicus* (formally known as *Clostridium glycolicum*), colonize leaf litter and mineral soil of oxygenated forest soils,[32, 34] suggesting that this organism can tolerate periods of increased O_2_ concentration. It is thus rather fitting to identify a *Cb*HydA1-like (likely, O_2_-tolerant) [FeFe] hydrogenase in this system. It is also intriguing that the second putative [FeFe] hydrogenase in *T. glycolicus* is missing the conserved proton-channeling Cys, a hallmark of the H_2_-producing [FeFe] hydrogenases. Therefore, it is more likely that the H_2_-production by this organism is due to *Tg*HydA1. Ultimately, this realization may point at *T. glycolicus* organism as a potential alternative H_2_-bioreactor host. However, further investigation is needed to uncover the physiological role of *Tg*HydA1 to take full advantage of this organism.

In conclusion, we have demonstrated that the ability to transition into the O_2_-stable state in [FeFe] hydrogenases shows a natural variation even between close homologues. Our study suggests that this deviation is controlled by the rate of enzyme reactivation, likely through the thermodynamic properties of the metallocofactors rather than structural mobility. We also show that this propensity for inactivation is accompanied by an increased stability upon O_2_ exposure. Therefore, the data presented in this work encourage closer examination of the energetics of electron transfer within the protein as a potentially important factor controlling O_2_ tolerance. Given that rapid reactivation is likely a key to designing efficient H_2_-producing systems, this work urges the development of a robust understanding of the structure-function relationships that dictate O_2_ tolerance in this class of enzymes.

## STATEMENTS AND DECLARATIONS

### Ethics declaration

not applicable.

### Consent to Participate

not applicable.

### Consent to Publish

not applicable.

## AUTHOR CONTRIBUTIONS

The manuscript was written through the contributions of all authors. All authors have given approval to the final version of the manuscript.

## ACKNOWLEDGMENTS

We greatfully thank Prof. John Golbeck for providing the cell line used in this work. This material is based upon work supported by the National Science Foundation under Award Number CHE-1943748 to A.S.

## ABBREVIATIONS

*E.coli*: Escherichia coli
*Tg*HydA1: [FeFe] hydrogenase 1 from *Terrisporobacter glycolicus*
*Cb*HydA1: [FeFe] hydrogenase 1 from *Clostridium beijerinckii*
PFV: Protein Film Voltametry
FTIR: Fourier Transform Infrared

## DATA AVAILABILITY

The authors declare that the data supporting the findings of this study are available within the paper and its Supplementary Information file. Should any raw data files be needed in another format, they are available from the corresponding author upon reasonable request.

## GRAPHICAL ABSTRACT

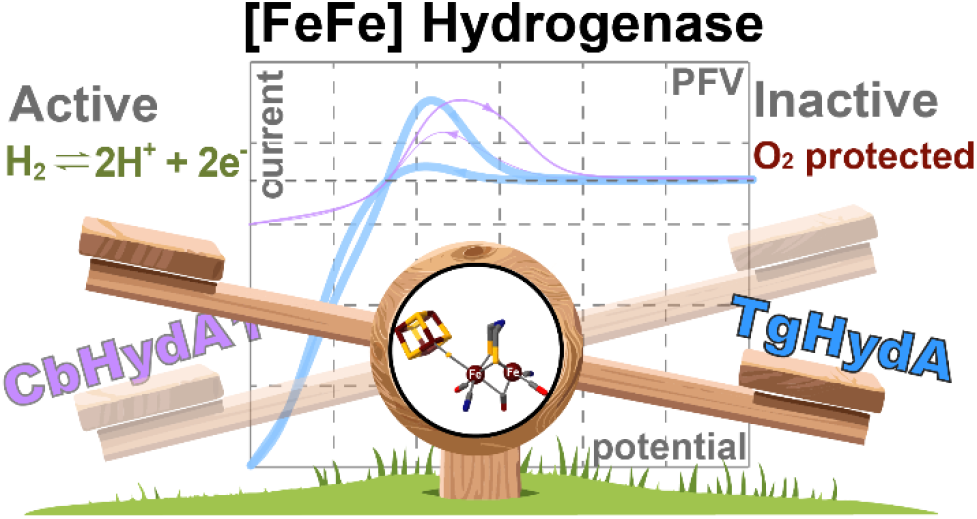

