## Supporting Information for "Redox Regulation in O_2_-Tolerant [FeFe] Hydrogenases: Insights from two homologues"

### Table of contents:

| | |
|---|---|
| Materials and Methods | 1 |
| Scheme S1 | 6 |
| Figure S1 | 7 |
| Table S1 | 8 |
| Figure S2 | 8 |
| Figure S3 | 9 |
| Figure S4 | 10 |
| Figure S5 | 11 |
| Figure S6 | 12 |
| References | 12 |

### Materials and Methods:

#### Cloning *Tg*HydA1

The complete gene of *Tg*HydA1 from *Terrisporobacter glycolicus* strain KPPR-9 (Locus tag: SAMN02910355_3663, NCBI Code: SFJ67572.1) was codon optimized for *Escherichia coli (E. coli).* The gene was augmented by an affinity tag with 6x His followed by a SUMO (GSDSEVNQEAKPEVKPEVKPETHINLKVSDGSSEIFFKIKKTTPLRRLMEAFAKRQGKEMDSLRFLYDG IRIQADQAPEDLDMEDNDIIEAHREQIGG) tag at the N-terminus. Strep-tag II sequence (WSHPQFEK) preceded by a spacer sequence (DIWSVGVKLFGGGSGGGSGGGS) was inserted at the C-terminus. The codon-optimized gene was inserted between Nco1 and Xho1 restriction sites in pET28a plasmid encoding for kanamycin resistance. The entire plasmid was acquired from Twist Bioscience. Plasmid containing the desired gene was then used to transform BL21 (DE3) ΔiscR *E. coli* cells (courtesy of the Golbeck laboratory at PSU) for recombinant expression.

Recombinant Expression

*Tg*HydA1 and *Cb*HydA1 were expressed per previously published protocol.[1] 150 ml LB starter cultures supplemented with 50 µg/mL kanamycin were grown at 37 ºC overnight, shaking at 200 rpm (New Brunswick Scientific C25 Incubator Shaker). Starter cultures were then transferred to 12 L LB media aliquoted in four 6 L flasks and grown aerobically, supplemented with 100 mM MOPS pH 7.4, 50 µg/mL kanamycin, 2 mM ferric ammonium citrate, and 2 mM L-cysteine hydrochloride, shaking at 200 rpm (New Brunswick Scientific C25 Incubator shaker). The cells were grown to an $OD_{600}$ of 0.6 ± 0.2. The culture was then allowed to cool on ice and supplemented with 0.5% glucose and 25 mM sodium fumarate. The culture was then transferred to six 2 L VWR bottles that were moved into an anaerobic glovebox (Coy Laboratories) under 97% $N_2$ and 3% $H_2$ and allowed to degas for 30 minutes. 750 µM IPTG and 750 µM NaDT were then added to the cells to induce protein expression overnight at 110 rpm at 18 ºC (Excella E25 Incubator Shaker). Cells were then harvested under strict anaerobic conditions by centrifugation in hermetically sealed 1 L centrifuge flasks at 4 ºC and 10,000 x g for 10 minutes (Thermo Fischer Scientific Sorvall$^{TM}$) and stored at -80 ºC until ready to be used.

Protein purification

All purification steps were carried out anaerobically inside a glovebox (Coy Labs) under 97% $N_2$ and 3% $H_2$ as per previously published protocol[1, 2]. The cells were resuspended in the lysis buffer containing 50 mM HEPES pH 7.5, 300 mM KCl, 5 mM β-mercaptoethanol and 5 mM imidazole supplemented with 1 mg/ml lysozyme, 0.1 mg/ml DNaseI, and 0.2 mg/ml phenylmethylsulphonyl fluoride and lysed by sonication (Sonics & Materials Inc. Model VCX 750) with one-second bursts at 80% amplitude with a nine-second delay between bursts for one hour stirring on ice. The lysate was then centrifuged at 60000 x g for 45 minutes to remove cell debris. The supernatant was then purified by Co-NTA resin using a gravity flow 30 ml column. The supernatant was loaded on the column and washed with 5 column volumes of wash buffer containing 50 mM HEPES pH 7.5, 300 mM KCl, 5 mM imidazole, and 10 mM ß-mercaptoethanol. The protein was then eluted with 300 mM imidazole buffer (50 mM HEPES pH 7.5, 300 mM KCl, 300 mM imidazole, 5 mM ß-mercaptoethanol). The eluent was transferred to a digestion buffer (50 mM HEPES pH 7.5, 300 mM KCl, 10% (v/v) glycerol, 5 mM DTT) using a PD-10 column (GE Life Sciences) after concentrating by centrifugation (Amicon). The protein was then incubated on ice overnight with ULP1 protease (obtained in-house with previously published protocols[3]) to remove the SUMO tag. The sample was then suspended in strep wash buffer (100 mM TAPS pH 8.0, 150 mM KCl, 2 mM sodium dithionite) and purified via Step-Tactin® Superflow high-capacity resin (IBA GmbH) with strep elution buffer (100 mM TAPS pH 8.0, 150 mM KCl, 2 mM sodium dithionite, and 2.5 mM desthiobiotin). The success of each step was verified by SDS-PAGE (see **Fig. S5**).

The iron-sulfur clusters of the protein were reconstituted by adopting protocols from Lanz et al.[4] The protein was resuspended in buffer containing 100 mM TAPS pH 8.0, 300 mM KCl, and 5mM dithiothreitol. The iron-sulfur clusters were then reconstituted by a slow addition of 15-fold excess $FeCl_3 \cdot 6H_2O$ over 30 minutes and 15-fold excess $Na_2S \cdot 9H_2O$ over 60 minutes. The protein was then allowed to incubate, stirring slowly on ice overnight (15 hours). The aggregates of unreacted iron and sulfur were pelleted by centrifugation at 10000 x g for 30 minutes at 4 ºC. The supernatant was then concentrated down by centrifugation using a 50 kDa Amicon centrifuge filter to 2.5 ml.

Synthetic reconstitution of the H-cluster was achieved as described previously.[1, 2, 5] In short, after reconstitution, the concentrated protein was buffer exchanged into 100mM TAPS pH 8.0. 150 mM KCl, and 5 mM sodium dithionite using a PD-10 column. $Fe_2[\mu\text{-}S_2C_2H_4NH](CO)_4(CN)_2$ precursor (Complex 1) was synthesized following previously published protocols[6–9], and the chemical identity was verified by FTIR. 5-fold excess Complex 1 was added to the protein and allowed to incubate, stirring slowly on ice overnight (15 hours). The excess of Complex 1 and sodium dithionite was removed, and the protein was simultaneously buffer exchanged into storage buffer (100 mM HEPES pH 7.5, 300 mM KCl, and 10% glycerol) by passing through a size-exclusion PD-10 column and stored at -80 ºC until ready to be used.

$O_2$ exposure and $H_2$ production assay

Sample solution of 1uM *Tg*HydA1 in 100 mM HEPES (pH 7.5), 150 mM KCl was repeatedly exposed to air. All samples (500 µL) were placed in 15 mL glass vials capped with a crimped rubber cap. Before any exposure of either control or aerobic samples, the protein was activated by pump-purging $H_2$ on a gas manifold inside the anaerobic glovebox (4 cycles, 30 sec pump, 30 sec purge) followed by incubation under 1 atm $H_2$ for additional 70 min. Subsequently, the samples to be exposed to air were taken out of the anaerobic chamber, fully uncapped and left on ice with a gentle stirring (<60 rpm) for 70-75 min. Anaerobic controls were also uncapped and placed on ice for 70 min, but inside anaerobic chamber (97%$N_2$, 3%$H_2$). For $H_2$ evolution assay, three 20uL aliquots of each sample solution were transferred into separate 15 mL glass vials (actual inner volume measured as 14.28 mL) already containing anaerobic solution of 480 ul 100 mM HEPES (pH 7.5) and 150 mM KCl for measurements in triplicates. The vials were capped and the headspace was flushed with ultra-pure He gas for 20 min on a gas manifold. The $H_2$ production was initiated by addition of 500uL of reduced methyl viologen solution (20 mM methyl viologen dissolved in 100 mM NaDT, 100 mM HEPES pH 7.5, 150 mM KCl). The headspace was analyzed for $H_2$ concentration with a Shimadzu GC-2010 Plus gas chromatograph equipped with an RT-Msieve 5A column (Restek) and a barrier ionization detector (Shimadzu BID 2010 Plus), using ultra-pure He as the carrier gas with a linear velocity of 30 cm/s with a split ratio of 50 or 100. The injection temperature was 150°C, the column temperature was 30°C, and the BID temperature was 280°C. Multiple measurements were taken in the course of 15 min. The sample volume of 100 uL was taken using 0.1 ml gas-tight syringe (Hamilton #1710).

FTIR

All FTIR spectra were collected on a Thermo Fisher Scientific Nicolet™50 FTIR instrument equipped with an $LN_2$-cooled MCT-A detector and a KBr beam splitter. The sample cell for room temperature measurements consisted of two 32 mm ODx3 mm $CaF_2$ windows (Crystran Ltd) spaced by a 40 µm PTFE spacer; two 32 mm OD, 15 mm ID silicone spacers of 1mm thickness sandwiched the $CaF_2$ windows and were placed in a custom aluminum holder, described previously.[2] The cryogenic measurements were conducted using a liquid He-flow cryostat (Oxford Instruments OptistatCF). The temperature was controlled by Oxford MercuryITC temperature controller. Cryogenic sample cell was composed of two 20 mm ODx2 mm $CaF_2$ windows (Crystran Ltd) with a

20 mm ODx15 mm IDx40 µm PTFE spacer between the windows. The windows were then placed in a custom copper holder designed according to the cryostat manufacturer's specifications. The spectra were collected over 2000 scans in the Single Beam mode. Room temperature measurements were taken at 2 $cm^{-1}$ resolution, while cryogenic measurements were taken at 1 $cm^{-1}$ resolution. Absorbance spectra were calculated from the single-beam trace using a single-beam background spectrum obtained with the identical buffer blank sample and under the same conditions. The water vapor peak subtraction was performed according to the previously described method,[2] followed by residual baseline modeling and subtraction using a cubic spline calculated from manually-selected data points. All data manipulations were performed using purpose-built scripts in MATLAB™. Processing scripts are available upon request.

Electrochemical experiments

The electrochemical cell contained a single junction Ag/AgCl reference electrode and a Pt counter electrode. The Ag/AgCl reference electrode was calibrated using methyl viologen as a standard ($E_0$ ($MV^{2+}$/MV•) = -446 mV vs NHE) prior to every measurement. The pyrolytic graphite electrodes were wet-polished with 1200-grit sandpaper (3M) and sonicated in deionized water for 10 minutes in a sonicator (VWR 50T). Background cyclic voltammogram and chronoamperogram traces for each electrode were collected using the same working electrode directly before loading the protein. Protein samples were diluted to 10 µM with 100 mM MES pH 6.5, 300 mM KCl. To generate protein films, 10 µl of the protein solution was pipetted onto the pyrolytic graphite electrode surface and allowed to adsorb onto the electrode for 5 minutes at room temperature. The excess liquid was removed from the electrode after the incubation period. All PFV experiments were carried out in buffers containing 300 mM KCl and appropriate buffers for the desired pH (100 mM MES pH 6.5, 100 mM HEPES pH 7.5, 100 mM TAPS pH 8.5). The buffer solutions were constantly purged with 100% $H_2$ during the measurements. The electrode rotation speed was set to 1000 rpm, and data were recorded at a scan rate of either 100 mV/sec or 10 mV/sec.

For chronoamperometry experiments, a cyclic voltammogram at 100 mV/s was recorded for each enzyme film before and after each potential step experiment to account for the film loss. The cell was set at $V_{red}$ = –800 mV vs NHE for 30 seconds to activate the enzyme before the beginning of the experiment, where the potential was switched between more oxidizing ($V_{inact}$) and more reducing ($V_{act}$) potentials. The experiment was run at each potential for 90 seconds.

Kinetic profile analysis was done using the following general derivations.

General kinetic model: $A \underset{k_{ba}}{\overset{k_{ab}}{\rightleftarrows}} B \underset{k_{cb}}{\overset{k_{bc}}{\rightleftarrows}} C$

In the current manuscript, we consider states A and B as active protein isoforms contributing to the catalytic current, while C is the inactive state.

A set of differential equations for the reaction above is:

$$\frac{dA}{dt} = -[A]k_{ab} + [B]k_{ba}$$

$$\frac{dB}{dt} = +[A]k_{ab} - [B]k_{ba} - [B]k_{bc} + [C]k_{cb}$$

$\frac{dC}{dt} = \qquad\qquad +[B]k_{bc} - [C]k_{cb}$

with $A(t) + B(t) + C(t) = const$

Matrix form of this set of equations: $\frac{d\mathbb{C}(t)}{dt} = \mathbb{C}(t)\mathbb{R}$ where $\mathbb{C}(t) = \begin{bmatrix} A(t) \\ B(t) \\ C(t) \end{bmatrix}$, $\mathbb{R} = \begin{bmatrix} -k_{ab} & +k_{ba} & 0 \\ +k_{ab} & -k_{ba} - k_{bc} & +k_{bc} \\ 0 & +k_{cb} & -k_{cb} \end{bmatrix}$

The solution has a form $\mathbb{C}(t) = e^{\mathbb{R}t}\mathbb{C}(0)$, where $\mathbb{C}(0) = \begin{bmatrix} A_0 \\ B_0 \\ C_0 \end{bmatrix}$ represents the initial (t=0) concentrations.

The values of the matrix exponential $e^{\mathbb{R}t}$ can be found by calculating eigenvectors and eigenvalues of $\mathbb{R}$, $\mathbb{R} = \boldsymbol{V}D\boldsymbol{V}^{-1}$, so that $e^{\mathbb{R}t} = \boldsymbol{V}e^{Dt}\boldsymbol{V}^{-1}$; where $D$ is a diagonal matrix containing eigenvalues (*1, $e_1$, $e_2$*).

The general solution can then be derived as:

$$A(t) = a_1(A_0 + B_0 + C_0) + \boldsymbol{e_1}b_1(+A_0x_2 + B_0y_2 - C_0z_2) + \boldsymbol{e_2}c_1(+A_0x_3 + B_0y_3 - C_0z_3)$$
$$B(t) = a_2(A_0 + B_0 + C_0) + \boldsymbol{e_1}b_2(-A_0x_2 - B_0y_2 + C_0z_2) + \boldsymbol{e_2}c_2(-A_0x_3 - B_0y_3 + C_0z_3)$$
$$C(t) = a_3(A_0 + B_0 + C_0) + \boldsymbol{e_1}b_3(-A_0x_2 - B_0y_2 + C_0z_2) + \boldsymbol{e_2}c_3(-A_0x_3 - B_0y_3 + C_0z_3)$$

where $e_1 = e^{-\frac{1}{2}(K_\Sigma + K_0)t}$, $e_2 = e^{-\frac{1}{2}(K_\Sigma - K_0)t}$

$a_1 = \frac{k_{ba}k_{cb}}{K_1}$, $\quad a_2 = \frac{k_{ab}k_{cb}}{K_1}$, $\quad a_3 = \frac{k_{ab}k_{bc}}{K_1}$

$b_1 = \frac{2(k_{bc}+k_{cb})-(K_0+K_\Sigma)}{4K_0K_1}$, $\quad b_2 = \frac{2k_{cb}-(K_0+K_\Sigma)}{4K_0K_1}$, $\quad b_3 = \frac{2k_{bc}}{4K_0K_1}$

$c_1 = \frac{2(k_{bc}+k_{cb})+(K_0-K_\Sigma)}{4K_0K_1}$, $\quad c_2 = \frac{2k_{cb}+(K_0-K_\Sigma)}{4K_0K_1}$, $\quad c_3 = \frac{2k_{bc}}{4K_0K_1}$

$\chi_2 = k_{ab}(K_0 - K_\Sigma)$, $\quad \chi_3 = k_{ab}(K_0 + K_\Sigma)$

$y_2 = k_{ab}(K_0 - K_\Sigma) + 2K_1$, $\quad y_3 = k_{ab}(K_0 + K_\Sigma) - 2K_1$

$z_2 = \frac{k_{cb}}{k_{bc}}((k_{ab} + k_{ba})(K_0 - K_\Sigma) + 2K_1)$, $\quad z_3 = \frac{k_{cb}}{k_{bc}}((k_{ab} + k_{ba})(K_0 + K_\Sigma) - 2K_1)$

$K_\Sigma = k_{ab} + k_{ba} + k_{bc} + k_{cb}$, $\quad K_1 = k_{ab}k_{bc} + k_{ab}k_{cb} + k_{ba}k_{cb}$,

$K_0 = \sqrt{(k_{ab} + k_{ba} - k_{bc} - k_{cb})^2 + 4k_{ba}k_{bc}}$

These equations were augmented by incorporating a mono-exponential decay due to gradual film loss. The decay constant was an additional fitting parameter, varied from one experiment to another. The catalytic current was assumed to be directly proportional to the sum of concentrations of the two active states with equal proportions.

$$i(t) = i_{max}(\mathrm{E})\big(A(t) + B(t)\big)e^{-t/t_{loss}}.$$

The $i_{max}$(E) was adjusted based on the later portions of the first potential step in either direction. Data manipulation and fitting were performed in MATLAB™ using purpose-built scripts that are available upon request. The fitting utilized Monte-Carlo-like random sampling of potential independent parameters in conjunction with a simplex fitting of the potential dependent parameters.

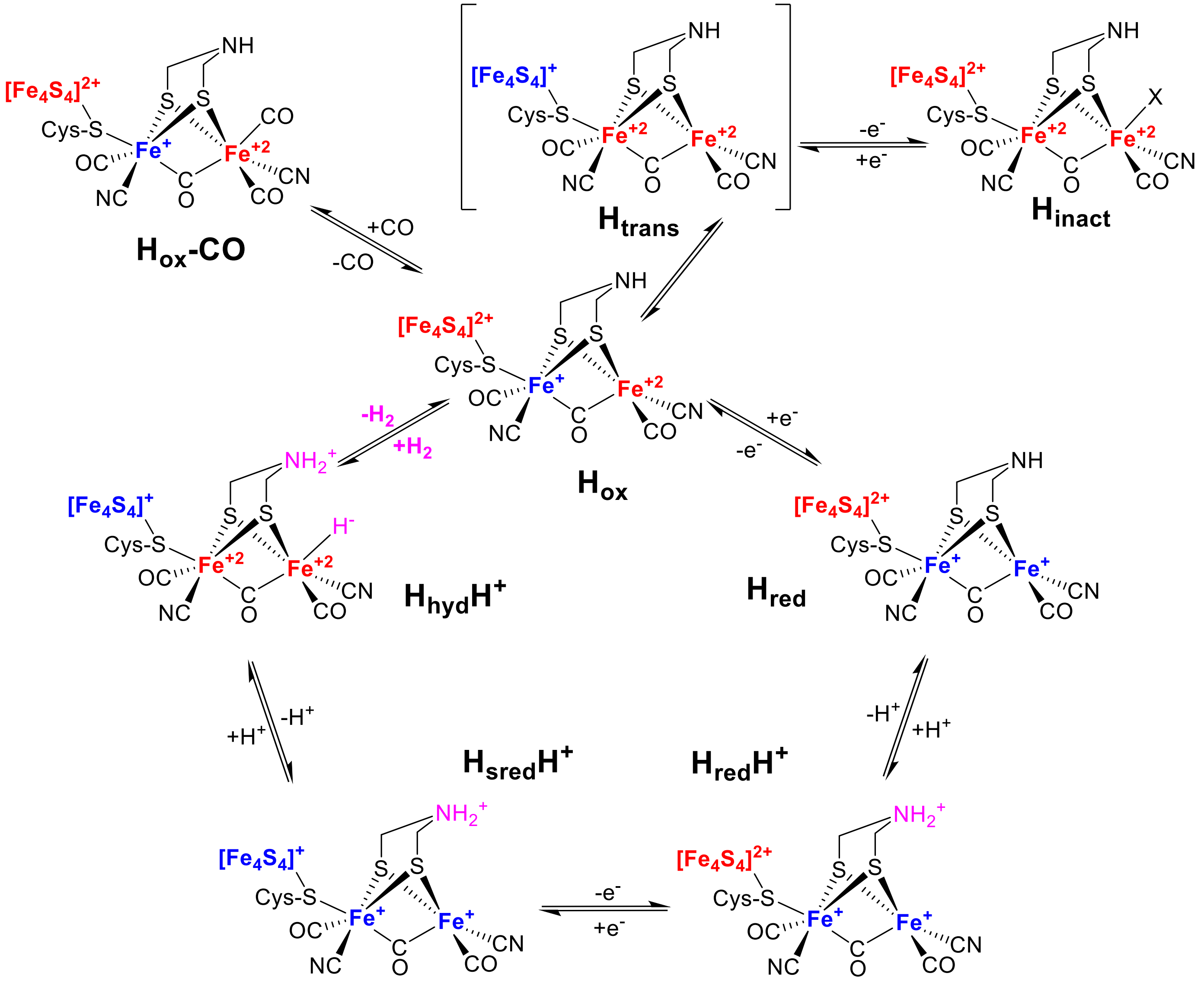


**Scheme S1.** Simplified proposed mechanism of $H_2$ evolution and $H_2$ oxidation by the H-cluster of [FeFe] hydrogenases, showing the relationship of states described in the main text. X in the $H_{inact}$ state designates $SH^-$ for DdHydAB and an unknown ligand for $O_2$-tolerant [FeFe] hydrogenases (*Cb*HydA1, *Tg*HydA).

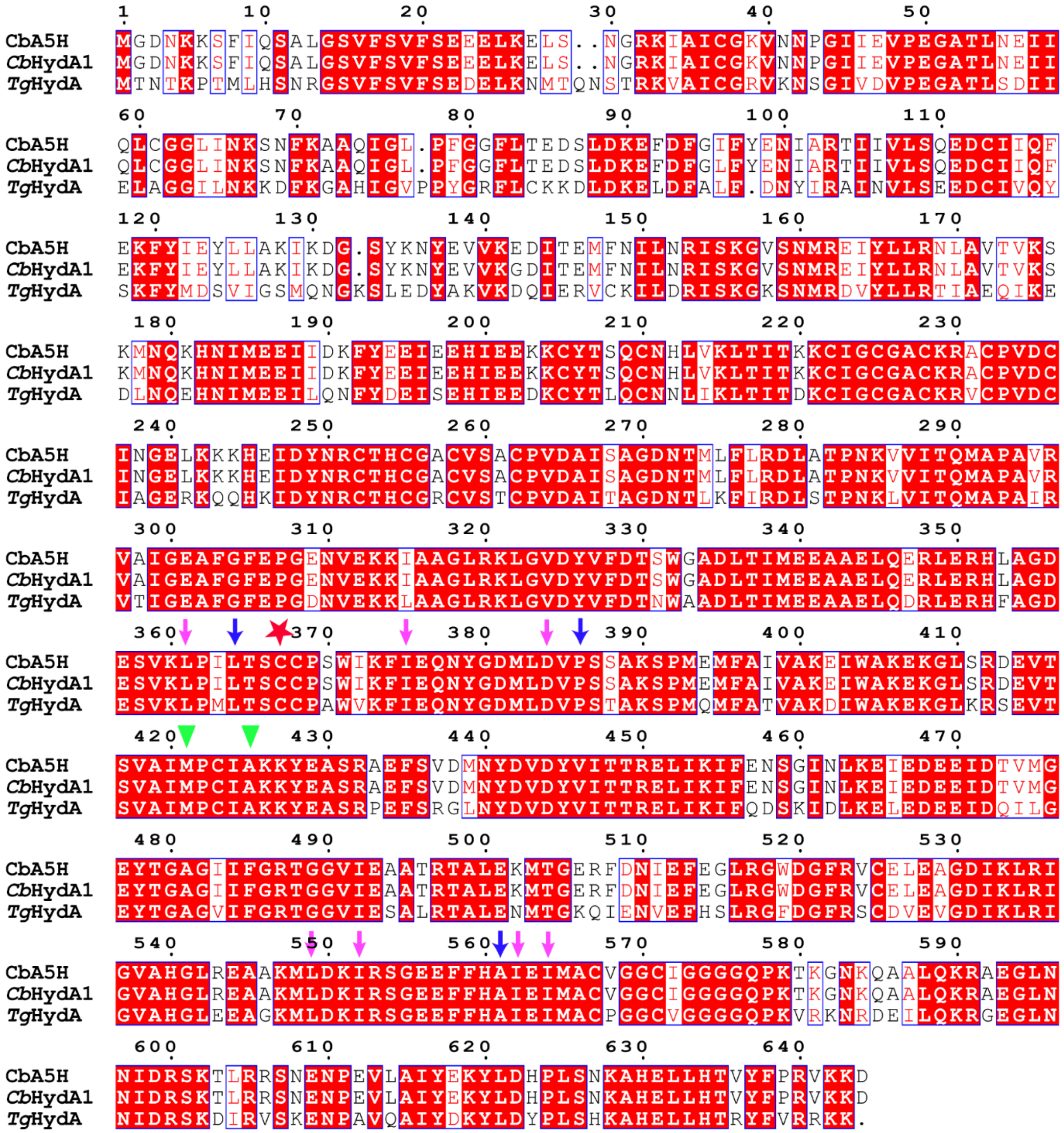


**Figure S1.** Sequence alignment of *Cb*A5H[10], *Cb*HydA1[1], and *Tg*HydA (NCBI Code: SFJ67572.1). Residues indicated by a red background and white text are highly conserved, while residues depicted in red text are partially conserved. Sequences were aligned using Clustal Omega[11]. The figure was prepared with ESPript[12]. The pink arrows indicate the amino acid residues that form the hydrophobic cluster centered around the proton-transporting cysteine-bearing loop per Ghosh et al.[13]. The blue arrows represent the amino acid residues that affect the TSC loop configuration and were mutated by Winkler et al.[14] The red star indicates Cys367, essential for catalytic activity. The green triangles indicate residues proposed to be involved in controlling catalytic bias based on crystallographic data of [FeFe] hydrogenase I from *Clostridium pasteurianum* (*Cp*I).[15] Percent identity and similarity between *Cb*A5H and *Cb*HydA1 is 99.69% and 99.84% respectively; *Cb*A5H and *Tg*HydA is 71.82% and 84.99% respectively; *Cb*HydA1 and *Tg*HydA is 72.06% and 84.86% respectively.

| State | Species | Terminal CN$^-$ | Terminal CO | Bridging CO | Ref |
|---|---|---|---|---|---|
| $H_{inact}$ | *Tg*HydA (RT) | 2107, 2080 | 2010, 1995 | 1838 | This work |
| | *Tg*HydA (15K) | 2111, 2081 | 2014, 1997 | 1838 | This work |
| | *Cb*HydA1 (RT) | 2107, 2080 | 2011, 2092 | 1840 | [1] |
| $H_{ox}$ | *Tg*HydA (RT) | 2089, 2054 | 1966, 1955 | 1782 | This work |
| | *Tg*HydA (15K) | 2091, 2054 | 1955, 1943 | 1762 | This work |
| | *Cb*HydA1 (RT) | 2092, 2081 | 1964, 1941 | 1800 | [1] |
| Unidentified | *Tg*HydA (RT) | 2099, 2071 | 1983, 1976 | 1825 | This work |
| | *Tg*HydA (15K) | 2099, 2069 | 1985, 1975 | 1830 | This work |
| $H_{trans}$ | *Dd*HydAB (RT) | 2100, 2075 | 1983, 1977 | 1836 | [16, 17] |

**Table S1.** Comparison of IR frequencies of CO and CN$^-$ bands of *Tg*HydA with previously published values. RT = room temperature.

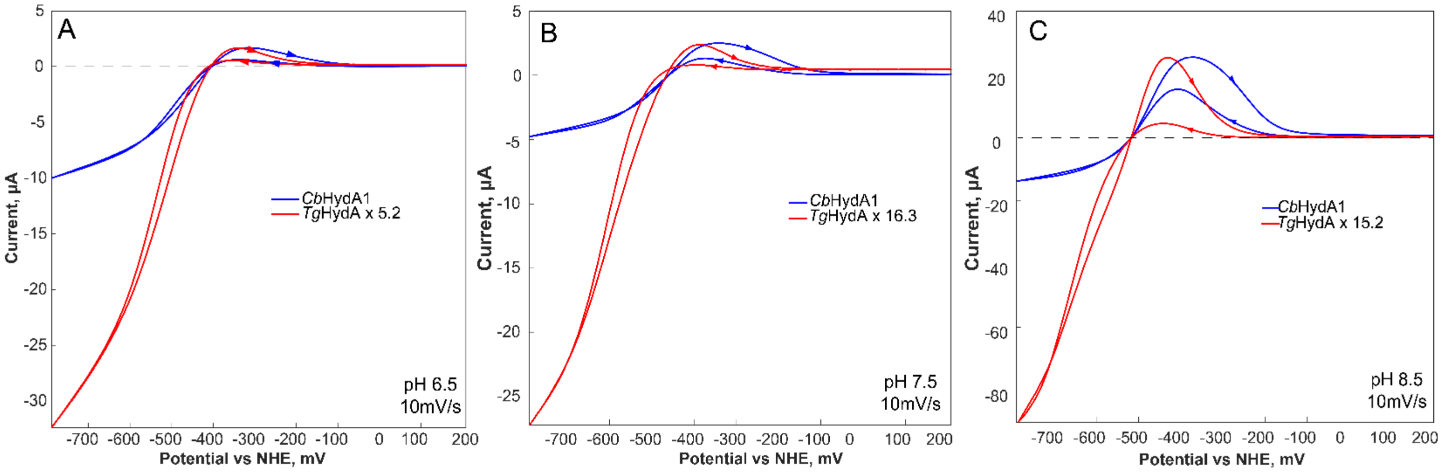


**Figure S2.** Comparison of protein film cyclic voltammetry traces of *Cb*HydA1 and *Tg*HydA performed at **A)** pH 6.5, **B)** pH 7.5 **C)** pH 8.5 at 10 mV/s scan rate. The trace of *Tg*HydA was scaled up by a factor of 5.2 in **A**, 16.3 in **B**, and 15.2 in **C** to match the maximum of the oxidizing current of *Cb*HydA1. Experiments were performed at room temperature under 100% $H_2$ atmosphere.

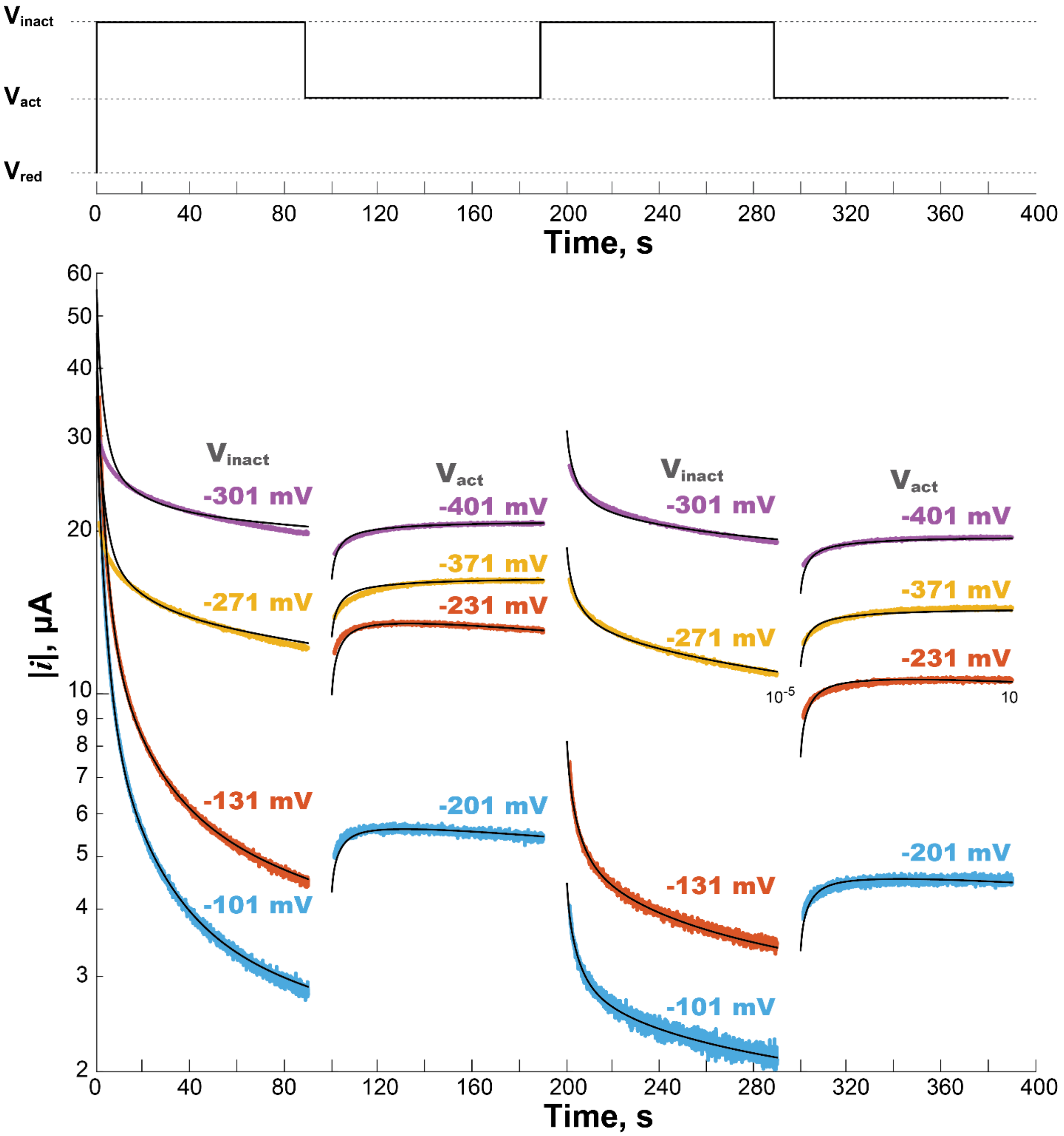


**Figure S3.** Chronoamperometry traces of *Cb*HydA1 obtained at pH 8.5 and room temperature. The general scheme of potential steps is shown in the top panel. $V_{red}$ = -800 mV vs. NHE in all experiments. In the bottom panel, colored chronoamperograms of the same color were obtained in a sequence within one experiment. Numbers above each trace indicate the potential, either $V_{inact}$ or $V_{act}$ (vs. NHE), at which the respective chronoamperogram was obtained. Black traces are fits according to equations and methods described in the Materials and Methods section above.

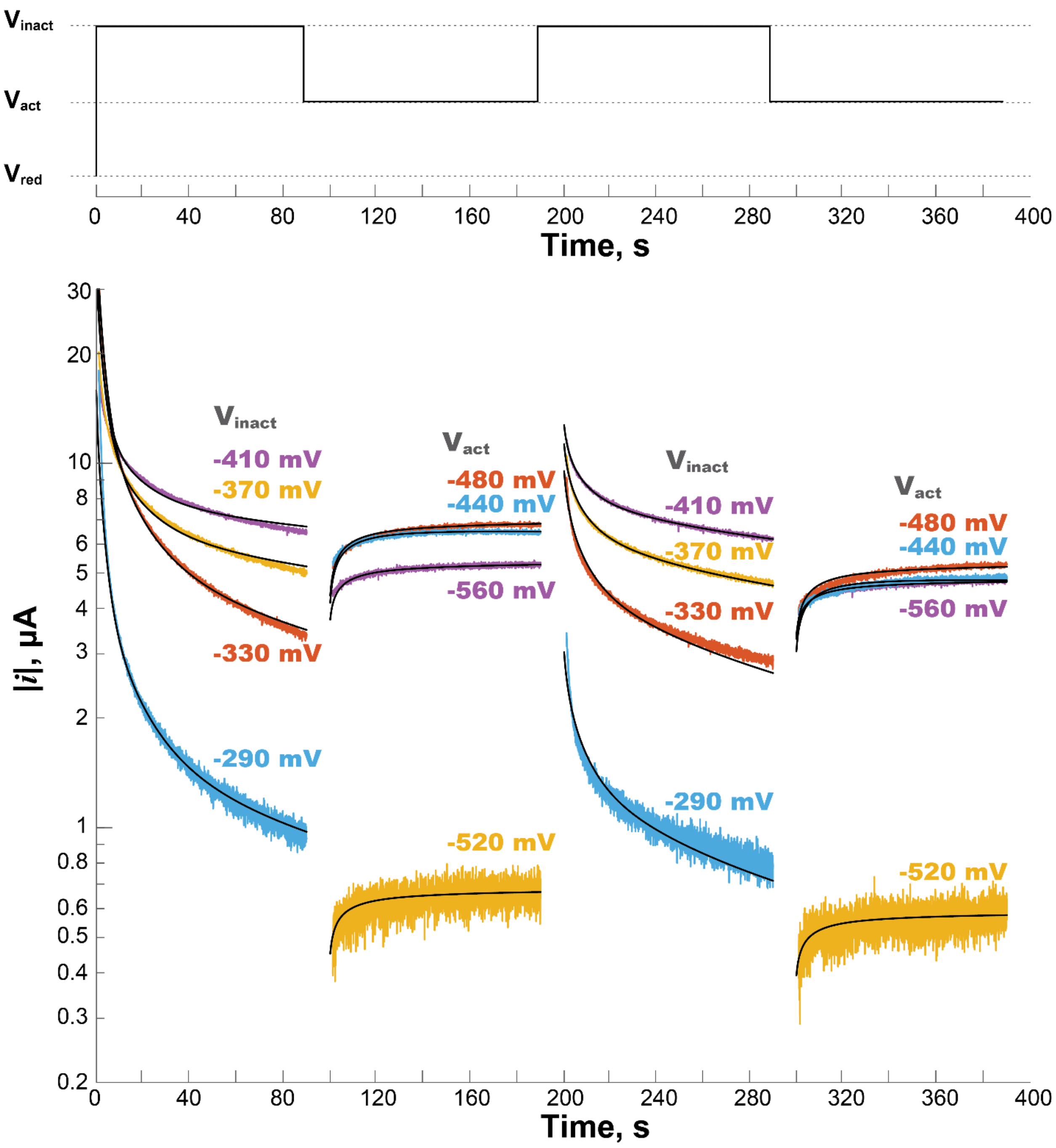


**Figure S4.** Chronoamperometry traces of *Tg*HydA obtained at pH 8.5 and room temperature. The general scheme of potential steps is shown in the top panel. $V_{red}$ = -800 mV vs. NHE in all experiments. In the bottom panel, colored chronoamperograms of the same color were obtained in a sequence within one experiment. Numbers above each trace indicate the potential, either $V_{inact}$ or $V_{act}$ (vs. NHE), at which the respective chronoamperogram was obtained. Black traces are fits according to equations and methods described in the Materials and Methods section above.

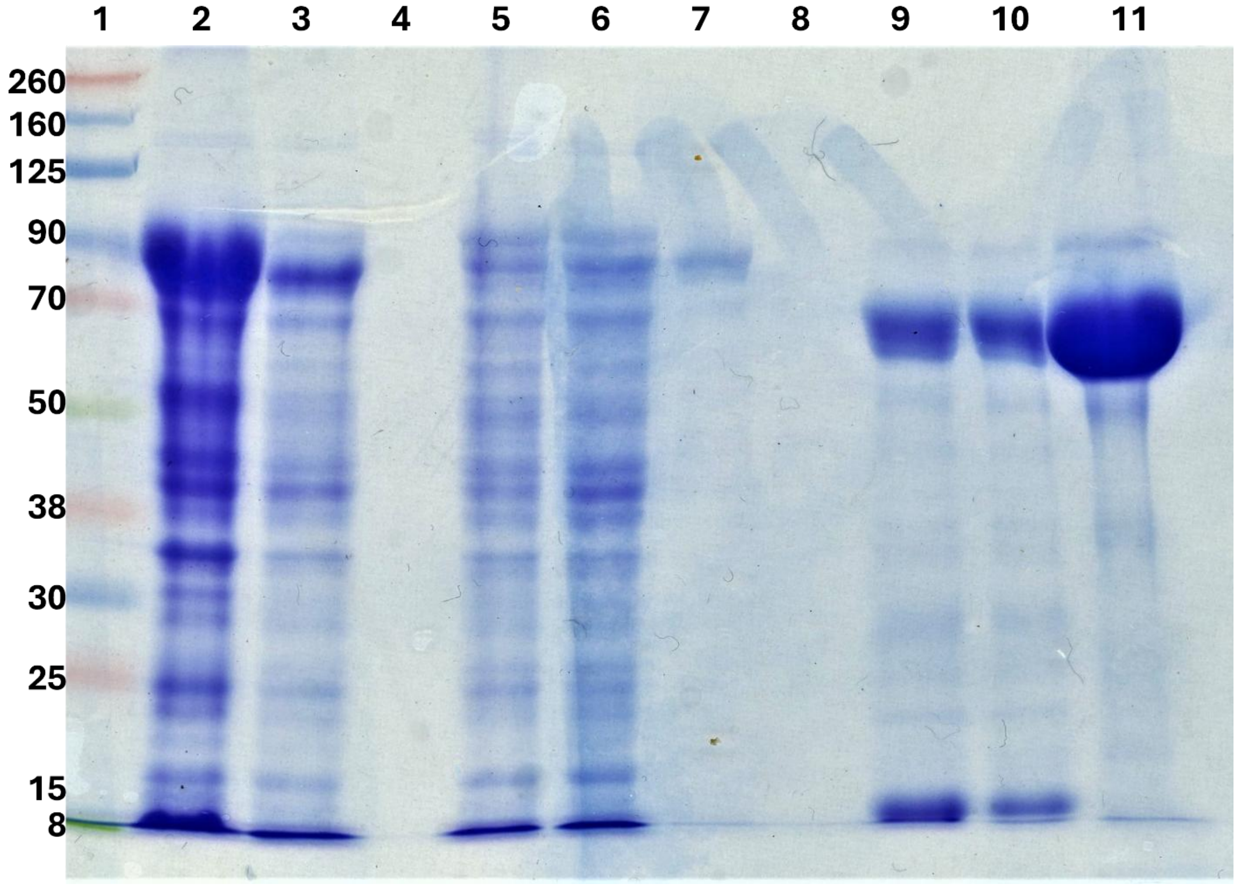


**Figure S5.** SDS-PAGE showing individual purification steps. Lane 1: ladder with molecular weights listed in kDa. Lane 2: pellet after homogenization, cell lysis, and centrifugation. Lane 3: supernatant after homogenization, cell lysis, and centrifugation. Lane 5: flow through from Co-NTA column. Lane 6: wash from Co-NTA column. Lane 7: Elution from Co-NTA column before cleaving the SUMO tag (Molecular weight of *Tg*HydA with SUMO Tag: 83kDa). Lane 9: Flowthrough from Strep-tactin column. Lane 10: Wash from Strep-tactin column. Lane 11: elution from Strep-tactin column after cleaving the SUMO tag with protease (Molecular weight of *Tg*HydA without SUMO tag: 72 kDa). Lanes 4 and 8 are blank.

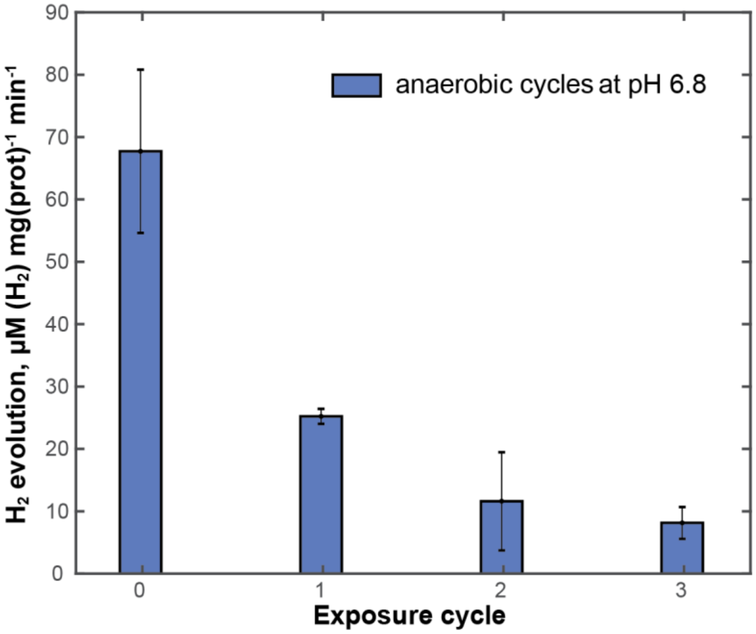


**Figure S6.** $H_2$ evolution activity of *Tg*HydA1 at pH 6.8 after repeated anaerobic treatment similar to the control experiment displayed on Figure 5 in the main text. Each cycle consisted of 75 min activation under 1 atm $H_2$ followed by incubation under 3%$H_2$ 97%$N_2$ on ice for additional 75min. Activity assays were performed by gas chromatography. Error bars are standard deviations calculated from measurements in triplicate.
